# Weak Membrane Interactions Allow Rheb to Activate mTORC1 Signaling Without Major Lysosome Enrichment

**DOI:** 10.1101/513473

**Authors:** Brittany Angarola, Shawn M. Ferguson

## Abstract

Stable localization of the Rheb GTPase to lysosomes is thought to be required for activation of mTORC1 signaling. However, the lysosome targeting mechanisms for Rheb remain unclear. We therefore investigated the relationship between Rheb subcellular localization and mTORC1 activation. Surprisingly, we found that Rheb was undetectable at lysosomes. Nonetheless, functional assays in knockout human cells revealed that farnesylation of the C-terminal CaaX motif on Rheb was essential for Rheb-dependent mTORC1 activation. Although farnesylated Rheb exhibits partial endoplasmic reticulum localization, constitutively targeting Rheb to ER membranes did not support mTORC1 activation. Further systematic analysis of Rheb lipidation revealed that weak, non-selective, membrane interactions support Rheb-dependent mTORC1 activation without the need for a specific lysosome targeting motif. Collectively, these results argue against stable interactions of Rheb with lysosomes and instead that transient membrane interactions optimally allow Rheb to activate mTORC1 signaling.

## Introduction

The mTOR complex 1 (mTORC1) signaling pathway plays a major role in matching cell growth and metabolism to ongoing changes in environmental conditions. Multiple signals converge on the surface of lysosomes to regulate the activity of the Rag and Rheb small GTPases that recruit and activate mTORC1 respectively. This has led to a widely accepted two step model for mTORC1 activation wherein Rags recruit mTORC1 to lysosomes followed by Rheb-dependent activation of mTORC1 kinase activity (Fig. S1A) (Ben-Sahra and Manning, 2017; Ferguson, 2015; Sancak et al., 2008; Saxton and Sabatini, 2017). However, even though farnesylation at its C-terminus has been widely accepted as a means of localizing Rheb to lysosomes, there is only limited direct support for this (Menon et al., 2014; Sancak et al., 2010; Sancak et al., 2008). Furthermore, as a farnesyl group is only expected to confer transient membrane interactions without selectivity for binding to lysosomes over other organelles (Silvius et al., 2006; Silvius and l’Heureux, 1994) the underlying mechanism of lysosome targeting is unexplained.

Adding to the confusion concerning the relationship between Rheb localization and function, it has also recently been proposed that Rheb instead resides on either the ER or Golgi and activates mTORC1 via contact sites between these organelles and lysosomes (Hao et al., 2018; Walton et al., 2018). These recent observations are paralleled by some older studies that also reported enrichment of over-expressed Rheb at the endoplasmic reticulum (Buerger et al., 2006; Hanker et al., 2010). The conflicting messages in these studies reveal uncertainty about both where Rheb functions within cells and how Rheb is targeted to its specific site of action.

To address these questions, we systematically investigated the relationship between Rheb localization and function in human cells. Surprisingly, our data indicates that Rheb does not require stable enrichment on a specific organelle. Instead, weak, non-selective, membrane interactions are sufficient to support mTORC1 activation. Collectively this data argues against the requirement for a stable and highly selective membrane interaction mechanism for optimal Rheb function. Our new data is not inconsistent with the widely accepted role for Rheb in activating mTORC1 at lysosomes. However, instead of stable residence of Rheb at lysosomes, we propose that transient membrane interactions satisfy the need to bring Rheb into proximity with mTORC1 that has been recruited to lysosomes via Rag-dependent mechanisms.

## Results and Discussion

### Rheb is not enriched on the surface of lysosomes

In contrast to the previously reported localization of Rheb to LAMP1-positive lysosomes in HeLa cells (Menon et al., 2014), we found that Rheb showed no major enrichment on such organelles (Fig. 1A), even though we used the same combination of antibody for Rheb detection and Rheb siRNA for verifying signal specificity (Fig. 1B). This difference in conclusions arises mainly from improved resolution of individual lysosomes which allowed for more stringent evaluation of colocalization. As Rheb is broadly found throughout cells, some Rheb inevitably overlaps with the LAMP1 signal. However, our analysis of the colocalization between Rheb and LAMP1 revealed that the degree of overlap between these two proteins was no greater than chance (Fig. 1C). As a control for the efficacy of the Rheb siRNA, immunoblotting experiments confirmed Rheb depletion following Rheb siRNA transfection and demonstrated that mTORC1 signaling was suppressed in these cells (Fig S1B-S1D).

**Figure 1.**
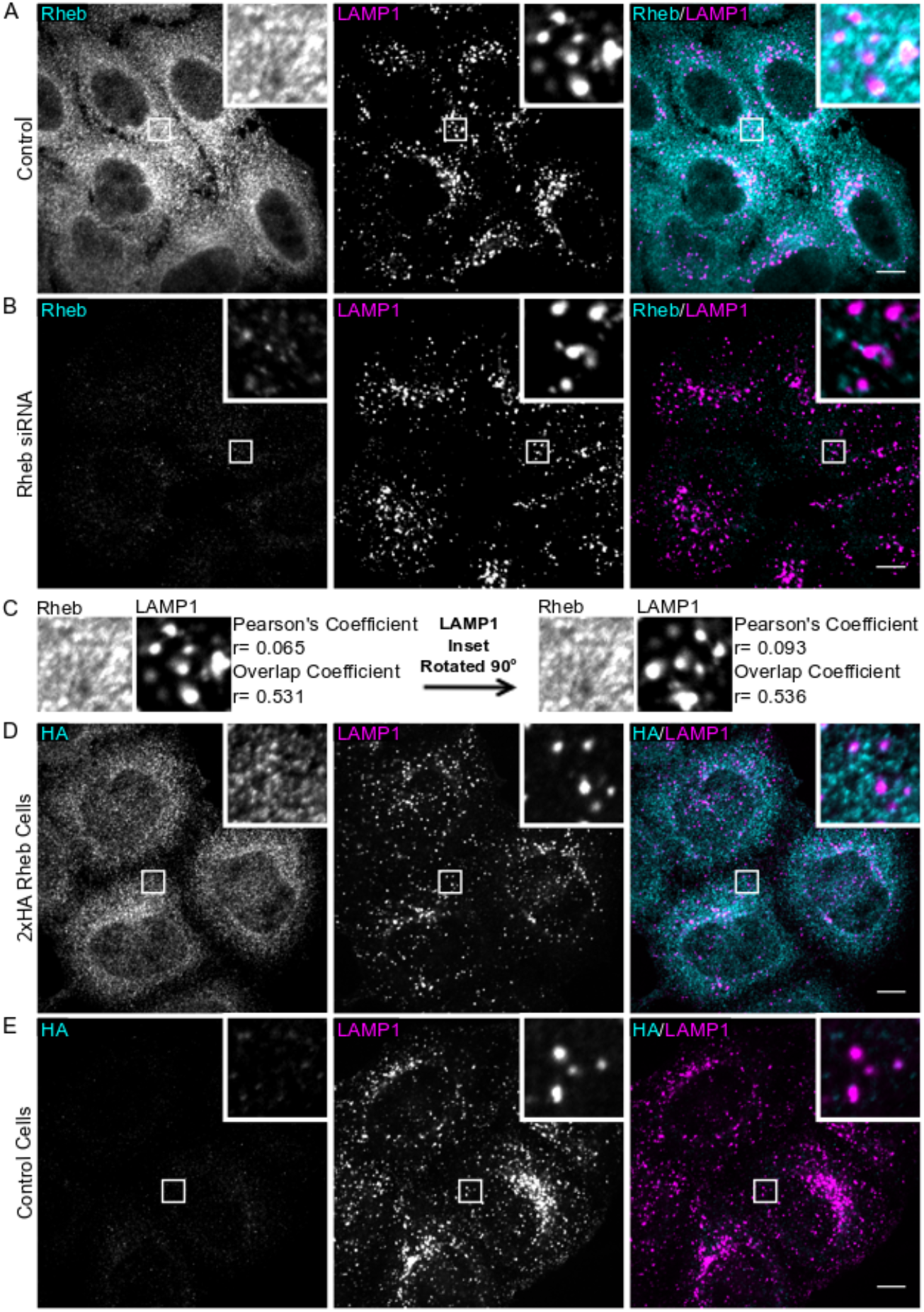
Rheb is not enriched on lysosomes. (**A and B**) Rheb and LAMP1 (late endosomes/lysosomes) immunofluorescence in HeLa cells that were transfected with control and Rheb siRNA respectively. **(C)** Correlation coefficients (ImageJ JACoP plugin) for Rheb and Lamp1 colocalization in the insets from panel A were calculated before and after rotating the LAMP1 image by 90°. (**D and E**) Immunofluorescence images of anti-HA and anti-LAMP1 staining in CRISPR-edited 2xHA-Rheb HeLa cells and control HeLa cells respectively. Scale bars, 10 μm.

As mTOR and many mTORC1 regulatory proteins exhibit dynamic changes in their levels at lysosomes in response to acute changes in amino acid availability [Fig. S1A; (Saxton and Sabatini, 2017)], we next examined the effect of amino acid starvation and re-feeding on Rheb localization. In contrast to mTOR, which was found throughout the cytosol under starved conditions but underwent a robust increase in its recruitment to lysosomes in response to amino acid re-feeding, Rheb localization was not responsive to such stimuli and failed to co-enrich with mTOR puncta even though mTORC1 signaling was stimulated in response to the amino acid re-feeding (Fig. S1E-S1G).

To generate an alternative tool for detecting the endogenous Rheb protein, we used CRISPR-Cas9 gene editing to insert a 2xHA epitope tag immediately downstream of the start codon in the endogenous Rheb locus in HeLa cells (Fig. S2A and S2B). However, the anti-HA immunofluorescence still did not show enrichment on lysosomes in these cells (Fig. 1D and 1E).

### Live cell imaging detects enrichment of GFP-Rheb at the endoplasmic reticulum

Preserving Rheb on lysosomes could require specialized fixation, permeabilization and/or antibody incubation conditions. Furthermore, our immunofluorescence experiments could have missed detecting a sub-population of Rheb at lysosomes due to epitope masking by interacting proteins. To circumvent such issues, we next examined the localization of GFP-tagged Rheb expressed at moderate levels in live HeLa cells and observed a combination of cytosolic and endoplasmic reticulum (ER)-like localization patterns (Fig. 2A). Rheb localization was further investigated in COS-7 cells as they contain a well-defined peripheral ER network that is highly suitable for live cell imaging studies (Fig. 2B) (Rowland et al., 2014). In addition to colocalizing extensively with mRFP-Sec61 (an ER protein), there was also a diffuse pool of Rheb in the cytosol (Fig. 2C). In contrast, GFP-Rheb and LAMP1-mCherry (lysosome marker) had distinct, non-overlapping, patterns of subcellular distribution. Interestingly, even though Rheb was not enriched on lysosomes, lysosomes were frequently adjacent to Rheb-positive ER tubules (Fig. 2D, Movie S1).

**Figure 2.**
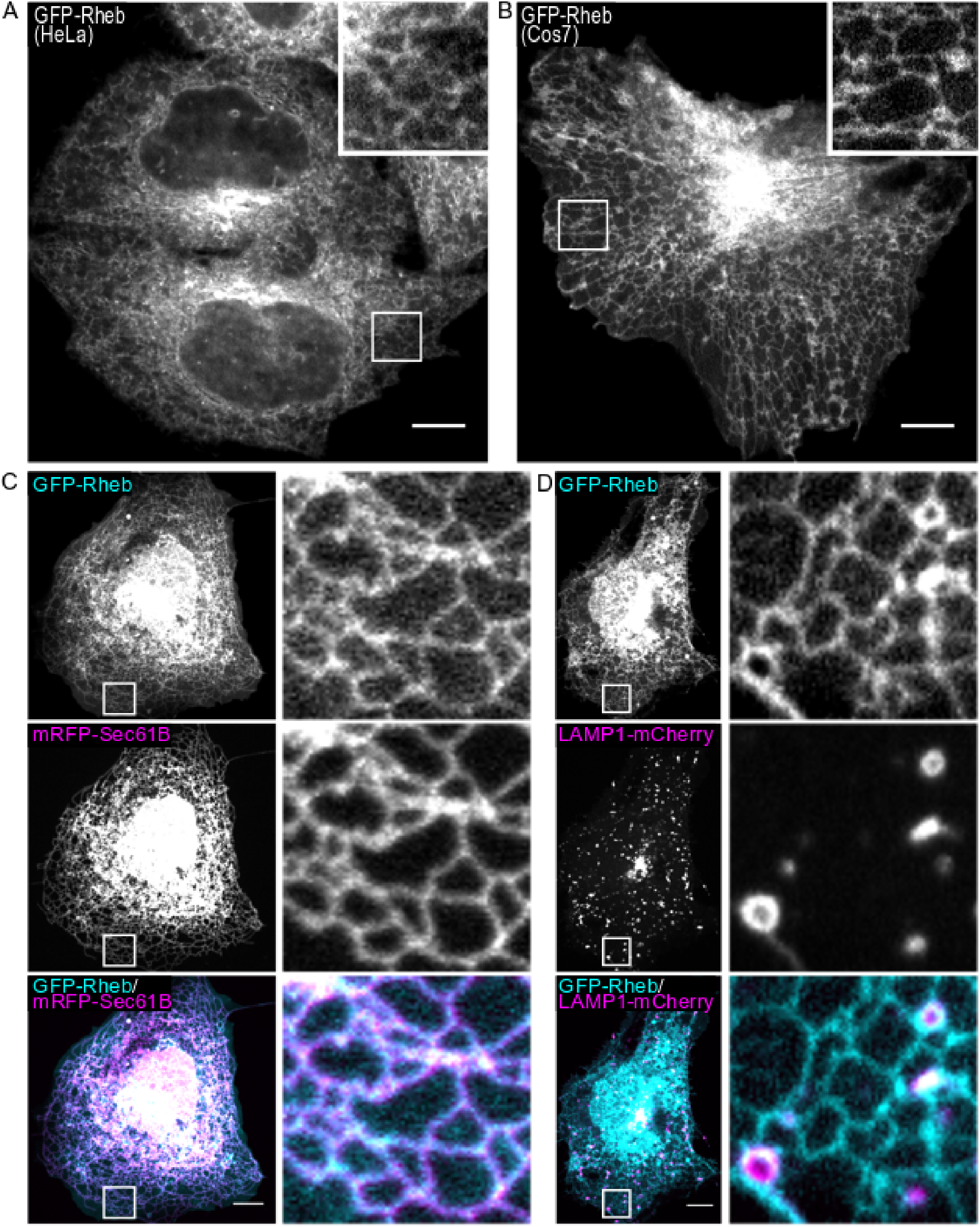
GFP-Rheb localizes to the ER and cytosol. (**A and B**) Spinning disk confocal live-cell imaging of GFP-Rheb in HeLa cells and COS-7 cells respectively. **(C)** GFP-Rheb and mRFP-Sec61B (ER marker) localization in COS-7 cells. **(D)** GFP-Rheb and LAMP1-mCherry (late endosomes and lysosomes) localization in COS-7 cells. Scale bars, 10 μm.

### Knockout cells reveal redundant functions for Rheb and RhebL1

In order to further investigate the relationship between Rheb sub-cellular localization and function, we next generated Rheb KO HeLa cells to use as a platform for measuring the ability of Rheb targeted to distinct subcellular locations to rescue mTORC1 signaling defects (Fig S2C). However, these Rheb KO cells maintained normal basal and serum-stimulated levels of mTORC1 signaling (Fig. S2C). This result indicated that Rheb is not absolutely essential for mTORC1 signaling and/or that cells can compensate for its absence. Consistent with the possibility of Rheb-independent mechanisms for mTORC1 activation, Rheb KO mouse fibroblasts were previously reported to exhibit reduced, but not eliminated, phosphorylation of mTORC1 targets (Goorden et al., 2011; Groenewoud et al., 2013).

We next considered a potential role for the Rheb-like 1 (RhebL1, also known as Rheb2) protein as a way for cells to activate mTORC1 signaling in the absence of Rheb. Few studies have focused on the RhebL1 protein. However, in addition to sharing sequence similarity with Rheb, RhebL1 over-expression was previously shown to stimulate mTORC1 signaling (Tee et al., 2005). We therefore used CRISPR-Cas9 genome editing to mutate both the Rheb and RhebL1 genes in Hela cells. After isolating clonal populations, we sought to identify cells harboring indels corresponding to frameshift mutations at the Cas9 target sites in both genes. The closest we came to achieving this goal was a cell line that contained one copy of Rheb with an in frame 6 base pair deletion and frameshift mutations in the remaining Rheb and RhebL1 alleles (Fig S2D, henceforth referred to as Rheb^edited^ cells). The absence of lines with frameshift mutations in all copies of both Rheb and RhebL1 likely reflects an essential role for Rheb/RhebL1 in supporting cell growth. Surprisingly, although Rheb protein levels were undetectable in the Rheb^Edited^ cells, basal levels of mTORC1 signaling were near normal (Fig. 3A and 3B). To test the possibility that traces of mutant Rheb protein containing the in frame mutation were responsible for the persistent mTORC1 signaling in the Rheb^Edlted^ cells, we further treated them with Rheb siRNA. This resulted in near complete suppression of basal mTORC1 signaling as assessed by measuring the phosphorylation state of multiple downstream targets (Fig. 3A and 3B; S2E and S2F). In light of these observations, we will subsequently refer to the Rheb^Edited^ cells that have been treated with Rheb siRNA as Rheb^Depleted^. These results indicate that Rheb and RhebL1 have redundant functions that are essential for mTORC1 signaling and provide an explanation for the previous reports of mTORC1 signaling in Rheb KO mouse embryonic fibroblasts (Goorden et al., 2011; Groenewoud et al., 2013).

**Figure 3.**
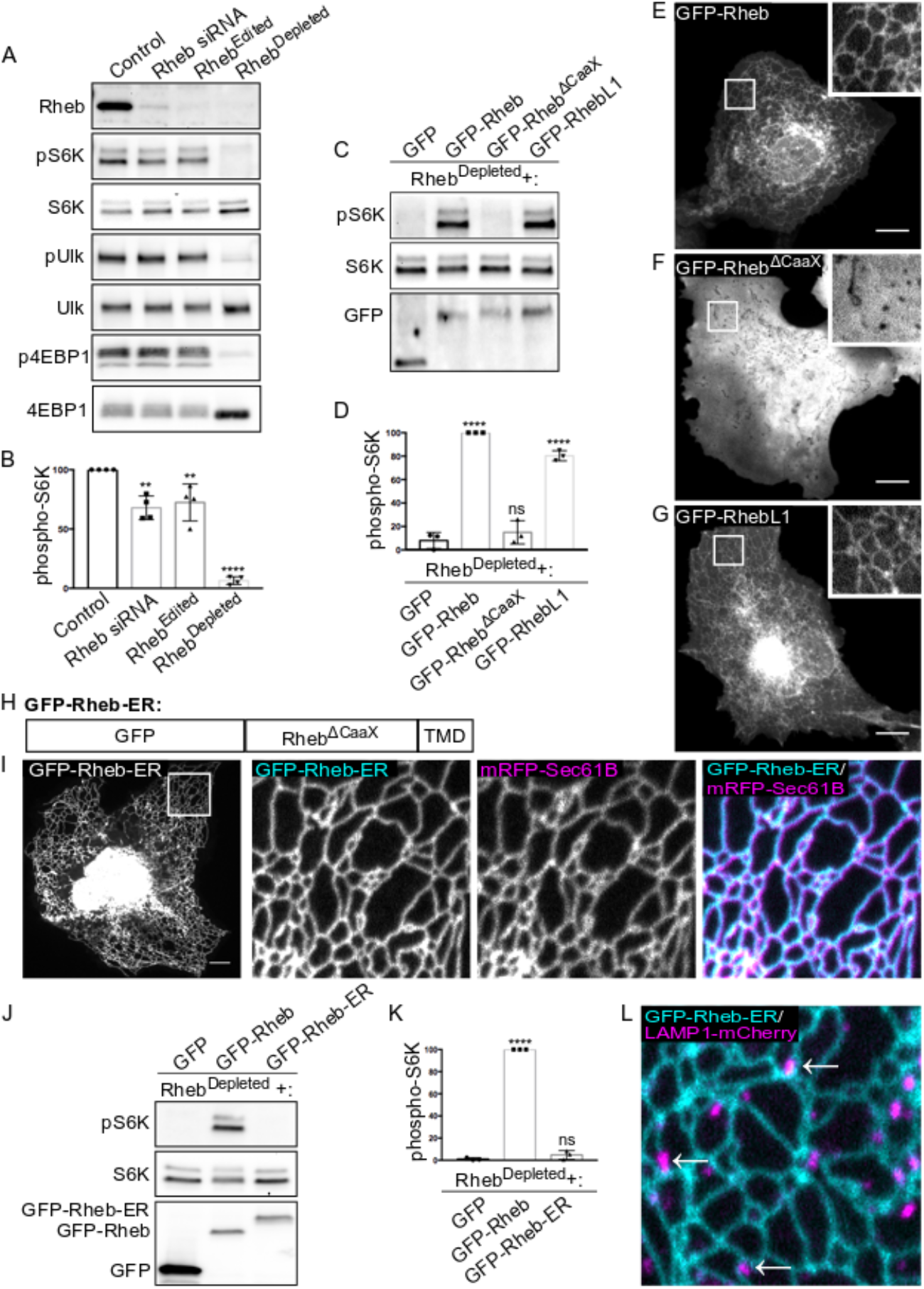
Development of Rheb^Depleted^ cells as a tool for testing relationships between Rheb localization and function. **(A)** Immunoblot analysis of Rheb levels and phosphorylation status of mTORCI substrates (S6K, ULK1, and 4EBP1) in Control, Rheb siRNA-treated, Rheb^Edited^ (Rheb hypomorph+RhebLI KO), and Rheb^Depleted^ (Rheb hypomorph+RhebL1 KO+Rheb siRNA) HeLa cells. **(B)** Quantification of phospho-S6K levels under the indicated conditions where phospho-S6K levels were divided by S6K and normalized to Control (**, P<0.01; ****, P<0.0001; ANOVA with Dunnett’s Multiple Comparisons Test, n=4). **(C)** Immunoblot analysis of phospho-S6K levels in Rheb^Depleted^ cells transfected with the indicated plasmids. **(D)** Quantification of phospho-S6K levels from panel C. The phospho-S6K levels were divided by S6K and GFP values to control for loading and transfection efficiency. Values were normalized to GFP-Rheb. Statistics were calculated in comparison to the GFP transfection. (****, P<0.0001; ANOVA with Dunnett’s Multiple Comparisons Test; n=3). **(E, F, G)** Live-cell images of GFP-Rheb, GFP-Rheb^ΔCaaX^, and GFP-RhebL1 in a COS-7 cells respectively. **(H)** Schematic of GFP-Rheb-ER chimera that contains N-terminal GFP, Rheb^ΔCaaX^, and the transmembrane domain of Cytochrome b5 (TMD). **(I)** Live-cell imaging of GFP-Rheb-ER localization. The leftmost image displays a low magnification view of GFP-Rheb-ER in a COS-7 cell. The three subsequent panels show higher magnifications of GFP-Rheb-ER and mRFP-Sec61B from the inset region. **(J)** Immunoblot analysis of phospho-S6K signaling in Rheb^Depleted^ cells transfected with the indicated plasmids. **(K)** Quantification of phospho-S6K levels in panel I. The phospho-S6K levels were divided by S6K and GFP values to control for loading and transfection. Values were normalized to GFP-Rheb. Statistics were calculated in comparison to GFP (****, P<0.0001; ANOVA with Dunnett’s Multiple Comparisons Test; n=3). **(L)** Image showing the spatial relationship between GFP-Rheb-ER on ER tubules and LAMP1-positive lysosomes. Scale bars, 10 μm.

### Rheb C-terminal farnesylation is essential for mTORC1 signaling and Rheb localization to the ER

Having established that mTORC1 signaling is eliminated in the Rheb^Depleted^ cells, we next used this new model system to investigate the subcellular targeting mechanisms that support Rheb function. mTORC1 signaling (S6K-T389 phosphorylation) was restored following expression of either GFP-Rheb or GFP-RhebL1 in Rheb^Depleted^ cells (Fig. 3C and 3D). Similar to GFP-Rheb, GFP-RhebL1 exhibited an ER-like localization pattern (Fig. 3E and 3G, S3A and B). Both Rheb and RhebL1 contain a C-terminal CaaX motif (cysteine followed by 2 aliphatic amino acids and X in the final position) wherein the cysteine is farnesylated (Fig. 4A; (Clark et al., 1997). A Rheb mutant that lacked the CaaX motif, was unable to rescue mTORC1 activity in Rheb^Depleted^ cells (Fig. 3C and 3D) and exhibited a diffuse subcellular distribution (Fig. 3F). These results indicate that Rheb cannot efficiently activate mTORC1 signaling from the cytosol and that farnesylation-dependent membrane interactions are essential for its function.

**Figure 4.**
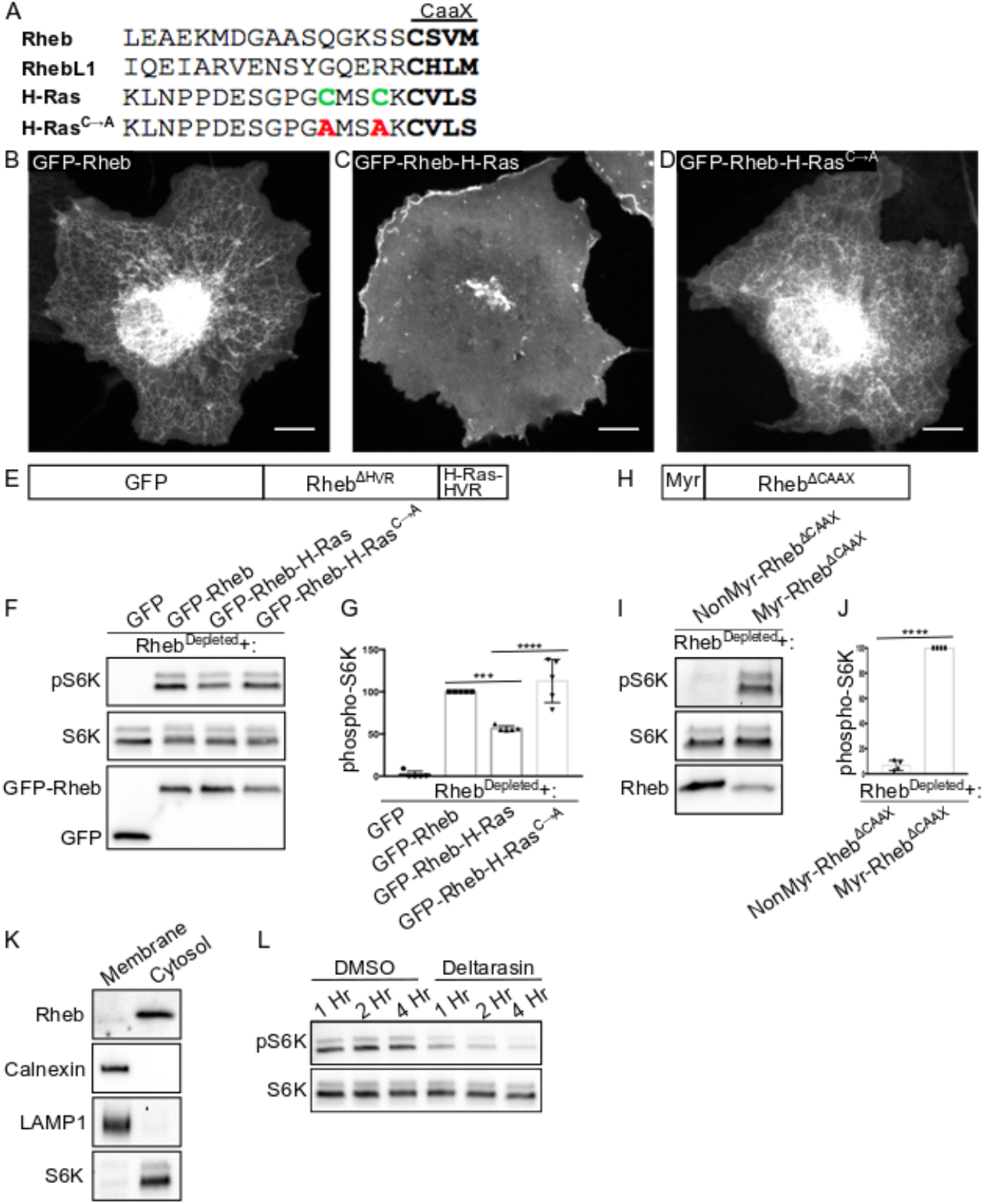
Weak membrane interactions are optimal for Rheb-dependent mTORC1 activation. **(A)** Alignment of the hypervariable regions of Rheb, RhebĲ and H-Ras proteins with CaaX motifs highlighted in bold and palmitoylated cysteines in green. These cysteines were mutated to alanines (red) in the GFP-Rheb-HRas^C➔A^ mutant. **(B-D)** Livecell images of GFP-Rheb, GFP-Rheb-HRas and GFP-Rheb-HRas^C➔A^ in COS-7 cells respectively. **(E)** Schematic of GFP-Rheb-H-Ras chimera that contains N-terminal GFP, Rheb^AHVR^, and the HVR of H-Ras. **(F)** Immunoblot analysis of phospho-S6K levels in Rheb^Depleted^ cells transfected with the indicated plasmids. **(G)** Quantification of phospho-S6K levels from panel F. Phospho-S6K levels were divided by S6K and GFP values to control for loading and transfection. Statistics were calculated in comparison to GFP-Rheb-HRas. Statistics were calculated in comparison to GFP-Rheb (***, P<0.001; ****, P<0.0001; ANOVA with Tukey’s Multiple Comparisons Test; n=5). **(H)** Schematic of myr-Rheb chimera that contains Rheb^ΔCaaX^ and a myristoylation consensus sequence. **(I)** Immunoblot analysis of phospho-S6K levels in Rheb^Depleted^ cells transfected with the indicated plasmids. **(J)** Quantification of phospho-S6K levels from panel I. Phospho-S6K levels were divided by S6K and Rheb values to control for loading and transfection (****, P<0.0001; Unpaired t test). **(K)** Immunoblot analysis of proteins identified in membrane and cytosolic fractions of HeLa cells. Scale bars, 10 μm. **(L)** Immunoblot analysis of phospho-S6K levels in HeLa cells treated with DMSO vehicle and 5 μM deltarasin for the times indicated.

Following the cytosolic farnesylation reaction, Rheb is further processed at the endoplasmic reticulum by Ras converting enzyme (RCE1) and isoprenylcysteine carboxymethyltransferase (ICMT) (Takahashi et al., 2005). This raised the possibility that the ER-localized pool of Rheb corresponds to newly synthesized Rheb protein that is undergoing these post-translational modifications. However, treatment of cells with cycloheximide to block new protein synthesis and thus allow any recently synthesized Rheb to complete its post-translational processing did not reduce Rheb abundance at the ER or result in the appearance of a lysosomal localization pattern (Fig. S4).

### Constitutive ER localization of Rheb does not support mTORC1 signaling

The prominent membrane contact sites between ER and lysosomes suggested that ER-localized Rheb might reach across such regions of proximity to activate mTORC1 that has been recruited to lysosomes via interactions with the Rags. Indeed, such a model was proposed in another recent study (Walton et al., 2018). To test this model, we generated a chimeric protein comprised of GFP-Rheb^ΔCaaX^ fused to the transmembrane domain of cytochrome B5 (Fig. 3H), a well-established ER targeting signal (Honsho et al., 1998). This chimeric protein localized to the ER (Fig. 3I) but did not stimulate mTORC1 activity in Rheb^Depleted^ cells (Fig. 3J and 3K) even though we still observed frequent contact between ER and lysosomes in cells expressing GFP-Rheb-ER (Fig. 3L).

### Rheb C-terminal farnesylation supports mTORC1 signaling without any requirement for additional targeting motifs in the surrounding hypervariable region

As C-terminal farnesylation is essential for the function of Rheb, we next focused on dissecting the role played by Rheb farnesylation and the possibility of additional regulation conferred by adjacent sequences in the Rheb C-terminal hypervariable region. In contrast to Rheb and RhebL1 whose ER and cytosolic localization is seemingly at odds with their ability to activate mTORC1 on lysosomes, other members of the Ras superfamily such as H-Ras localize robustly to their major site of action at the plasma membrane (Choy et al., 1999; Hancock et al., 1990). The plasma membrane localization of H-Ras depends on C-terminal farnesylation accompanied by 2 additional cysteine residues within the adjacent hypervariable region that are palmitoylated (Fig. 4A) (Choy et al., 1999; Hancock et al., 1990). It was also reported that the last 15 amino acids of Rheb acts as a lysosome targeting signal (Sancak et al., 2010). This suggested that this sequence should contain an additional determinant that cooperates with farnesylation to target Rheb to lysosomes.

To investigate the role of the CaaX motif and its flanking sequences in determining the subcellular localization and function of Rheb, we generated a chimera wherein the Rheb C-terminus was replaced with the last 20 amino acids from H-Ras (Fig. 4E). This Rheb-H-Ras protein localized predominantly to the plasma membrane (Fig. 4B and 4C) and was less effective than full length Rheb at activating mTORC1 signaling (Fig. 4F and 4G). Although, the predominant localization of H-Ras to the plasma membrane is seemingly at odds with the ability of the Rheb-H-Ras chimera to moderately activate mTORC1 signaling, the H-Ras C-terminus undergoes cycles of depalmitoylation that allow it to transiently visit intracellular membranes (Rocks et al., 2005). To test the idea that farnesylation alone is the key determinant of Rheb localization and function, we generated a mutant Rheb-H-Ras chimera that lacks the palmitoylated cysteines in the hypervariable region (Fig. 4A). This protein was no longer enriched at the plasma membrane and had a similar localization pattern to the WT Rheb (Fig. 4B and 4D). It also promoted mTORC1 activity just as well as the WT GFP-Rheb (Fig. 4E and 4F). As Rheb, RhebL1 and the palmitoylation deficient H-Ras hypervariable regions lack sequence similarity beyond their CaaX motifs (Fig. 4A), these results indicate that farnesylation, independent from any other major determinants within the Rheb hypervariable region, is both necessary and sufficient for the ability of Rheb to interact with membranes and support mTORC1 signaling.

### N-terminal myristoylation can substitute for Rheb C-terminal farnesylation

Our results challenge expectations that Rheb stably resides at lysosomes via a specific targeting signal that resides within its C-terminus. To further test this conclusion, we sought an alternative method for dynamically targeting Rheb to membranes. For this purpose, we generated a Rheb mutant that lacks the C-terminal CaaX motif and therefore cannot be farnesylated but instead contains a myristoylation signal at its N-terminus (Fig. 4H). This strategy was based on the following logic: 1) Myristoylation, like farnesylation supports only transient membrane interactions; 2) N-terminal myristoylation was previously shown to be an effective substitute for C-terminal farnesylation in H-Ras (Cadwallader et al., 1994); and 3) Recent phylogenetic analysis identified Rheb genes in some species that lack C-terminal farnesylation but are predicted to be myristoylated at their N-termini (Zahonova et al., 2018). Interestingly, although a Rheb mutant that is neither farnesylated or myrisotylated failed to promote mTORC1 signaling in Rheb^Depleted^ cells, myristoylated Rheb stimulated S6K phosphorylation (Fig. 4I and 4J). The functionality of myristoylated Rheb indicates that a weak, non-selective, membrane interaction is the minimal targeting signal that is required for Rheb function. This observation also argues against the possibility that farnesylation is essential for other aspects of Rheb function such as its ability to activate mTORC1. This interpretation is further corroborated by in vitro experiments that ruled out a direct requirement for Rheb farnesylation in mTORC1 activation (Sato et al., 2009). Consistent with a model wherein Rheb C-terminal farnesylation only supports transient membrane interactions, subcellular fractionation revealed that Rheb was predominantly found in the cytosolic fraction and only minimally present in the membrane fraction (Fig. 4K).

Dynamic redistribution of Rheb between membranes is regulated by its interaction with PDEδ (Ismail et al., 2011; Kovacevic et al., 2018). The binding of farnesylated proteins to PDEδ can be blocked by drugs such as deltarasin, that compete for binding to the hydrophobic cavity of PDEδ (Zimmermann et al., 2013). Consistent with previous reports (Papke et al., 2016; Zimmermann et al., 2013), we observed that mTORC1 signaling was suppressed in deltarasin-treated cells (Fig. 4L). Although this could reflect effects on multiple farnesylated proteins beyond Rheb, this result is nonetheless consistent with a role for dynamic, PDEδ-dependent, redistribution of Rheb between subcellular membranes in supporting mTORC1 signaling.

### Implications of weak Rheb membrane interactions for mTORC1 signaling

The minimal membrane targeting affinity and specificity of Rheb/RhebL1 contrasts with other members of the Ras superfamily that exhibit robust targeting to the membranes where they recruit their downstream effectors. However, Rheb is distinct from other members of the Ras family in at least two important ways. First, Rheb is not required for mTOR recruitment to the surface of lysosomes (Fig. S2G). mTORC1 localization to lysosomes is instead dependent on interactions between the Raptor subunit of mTORC1 and the Rag GTPases (Sancak et al., 2010). Second, recent structures of Rheb-mTORC1 complexes revealed that Rheb stimulates mTORC1 catalytic activity by binding to mTOR and inducing a conformational change that rearranges the mTOR kinase domain (Yang et al., 2017). Once this activation takes place, mTORC1 phosphorylates diverse substrates that reside throughout the cell (Ben-Sahra and Manning, 2017; Gonzalez and Hall, 2017; Saxton and Sabatini, 2017). Thus, the modest membrane affinity that is provided by Rheb farnesylation may represent an optimal solution for reducing dimensionality to facilitate interactions with lysosome-localized mTORC1 while also allowing the activated Rheb-mTORC1 complex to subsequently leave the lysosome to phosphorylate downstream targets that reside elsewhere in the cell.

### Evidence in favor of dynamic interactions between Rheb and endo-lysosomal membranes as the key determinant mTORC1 activation

A predominantly ER and cytosolic steady state localization of the Rheb protein challenges the idea that Rheb stably resides on lysosomes while waiting for the opportunity to activate mTORC1. Nonetheless, there are several arguments in support of a function of Rheb at lysosomes that could be mediated by transient interactions. First, considerable evidence indicates that Rag-dependent recruitment of mTORC1 to lysosomes is a prerequisite for mTORC1 activation and that such activation also depends on Rheb (Bar-Peled et al., 2013; Bar-Peled et al., 2012; Betz and Hall, 2013; Lim and Zoncu, 2016; Petit et al., 2013; Sancak et al., 2010; Sancak et al., 2008; Tsun et al., 2013; Wolfson et al., 2017). Second, TSC, the Rheb GAP is found at lysosomes, particularly in nutrient starved cells [Fig. S1; (Carroll et al., 2016; Demetriades et al., 2014; Demetriades et al., 2016; Menon et al., 2014)]. Third, constitutively targeting TSC to lysosomes suppresses mTORC1 signaling (Menon et al., 2014). Fourth, although the Rheb protein in most commonly studied model organisms is targeted to membranes via farnesylation of a C-terminal CaaX motif, a recent phylogenetic analysis of Rheb protein sequences revealed that Rheb homologs in some species lack lipidation and are instead predicted to interact with membranes via an N-terminal FYVE domain (Zahonova et al., 2018). In others, the C-terminal CaaX motif is accompanied by an N-terminal PX domain (Zahonova et al., 2018). Although these bioinformatic predictions remain to be experimentally validated, PX and FYVE domains commonly recruit proteins to endosomal membranes via their ability to bind to phosphatidylinositol-3-phosphate (PI3P) with modest affinity (Burd and Emr, 1998)(Gaullier et al., 1998). This evolutionary selection for Rheb targeting mechanisms that involve non-covalent interactions with endosomal lipids is consistent with a conserved function for Rheb in the endolysosomal pathway.

As an alternative to the possibility that Rheb dynamically visits the surface of lysosomes in order to activate mTORC1, it was recently reported that Rheb instead resides on the Golgi and activates mTORC1 via contact sites between the Golgi and lysosomes (Hao et al., 2018). However, this study relied on assessing the ability of heavily over-expressed, Golgi-targeted Rheb, to hyper-activate mTORC1 signaling. Furthermore, the Golgi targeting strategy was based on engineering an artificial interaction between Rheb and Rab1a. Although this resulted in significant localization of Rheb to the Golgi and Rheb-dependent activation of mTORC1 signaling, Rab1a is a prenylated protein that cycles on and off of Golgi membranes (Smeland et al., 1994). Given our results with the predominantly plasma membrane localized Rheb-H-Ras chimera (Figure 4) as well as our observations that only very low levels of Rheb are required for mTORC1 activation (Figure 3), caution must be applied with interpreting how the steady state localization of an overexpressed Rheb protein that undergoes dynamic cycling on and off membrane relates to the actual site of mTORC1 activation.

### Potential implications of Rheb localization to the ER

It was recently proposed that moving lysosomes towards the cell periphery suppresses mTORC1 activity by limiting their proximity with ER-localized Rheb (Walton et al., 2018). Although this idea is intriguing, we observed that even peripheral lysosomes still maintain contact with Rheb-positive ER tubules (Fig. 3, Movie S1). Furthermore, the inability of constitutively ER-localized Rheb to activate mTORC1 in spite of maintained ER-lysosome contact sites argues against a model wherein ER-localized Rheb activates mTORC1 by reaching across such contact sites (Fig. 3). Finally, it has long been known that the farnesylated CaaX motif is sufficient to target proteins such as GFP to the ER (Choy et al., 1999). Therefore, Rheb localization to the ER may simply reflect the default localization for a farnesylated protein that lacks other strong subcellular localization signals.

### Conclusions

Although considerable evidence indicates that lysosomes are a major intracellular site where nutrient and growth factor signals are integrated in order to match mTORC1 signaling to ongoing environmental changes, our new observations argue against the presence of a large steady state pool of Rheb at lysosomes. Nor did our results support the existence of a selective lysosome targeting signal within the Rheb hypervariable region. Instead, we propose that transient, farnesylation-dependent, membrane interactions have been selected by evolution as optimal for the Rheb-mediated activation of mTORC1 at lysosomes. By overturning widely held beliefs concerning stable residence of Rheb at lysosomes, our new findings will guide future studies that focus on understanding how dynamic Rheb membrane interactions are coordinated with the Rag-dependent localization of mTORC1 to lysosomes to activate mTORC1 signaling in health and disease.

## Acknowledgements

This research was supported in part by grants from the NIH (GM105718) and the Ellison Medical Foundation to SMF. BA was supported by an NSF Graduate Research Fellowship (DGE1752134). Pamela Torola contributed to early phases of this project and Arun Tharkeshwar provided helpful advice related to cell fractionation. Microscopy studies were supported by the Yale University Program in Cellular Neuroscience, Neurodegeneration and Repair imaging facility.

## Methods

### Cell Culture and Transfection

HeLa M cells and COS7 cells (both provided by P. De Camilli, Yale University, New Haven) were grown in high-glucose Dulbecco’s Modified Eagle’s medium (DMEM) with L-glutamine, 10% fetal bovine serum, and 1% penicillin/streptomycin supplement (ThermoFisher Scientific, Waltham, MA and Corning, Corning, NY). To starve cells of amino acids, cells were incubated in amino acid-free RPMI media (US Biologicals, Swampscott, MA) for 2 hours. Cells were re-fed with RPMI containing 1X MEM amino acid supplement (Invitrogen) for 20 minutes. To starve cells of growth factors, cells were incubated in serum-free DMEM overnight. Cells were re-fed for 30 minutes with complete media.

To perform plasmid transfections, 500 ng of plasmid DNA, 1.5 μl of Fugene 6 transfection reagent (Promega, Madison, WI) and 100 μl of OptiMEM (ThermoFisher Scientific) were added to 80,000 cells per well in a 6-well dish. For siRNA transfections, 5 μl of RNAiMAX (ThermoFisher Scientific), 500 μl of OptiMEM and 5 μl of 20 μM siRNA stock were added to a subconfluent dish of cells (80,000 cells per well in a 6-well dish). Cells were incubated for 48 hours post-transfection prior to experiments. Control siRNA (5’-CGUUAAUCGCGUAUAAUACGCGUAT-3’) was purchased from Integrated DNA Technologies (IDT, Coralville, IA) and a previously described Rheb siRNA (Menon et al., 2014) was purchased from Cell Signaling Technology (#14267, CST, Danvers, MA). Drugs utilized in this study: Deltarasin (Cayman Chemical, Ann Arbor, MI, #9001536) and Cycloheximide (Millipore Sigma, Burlington, MA, #239765).

### Plasmids

pRK5 plasmid encoding HA-GST human Rheb was acquired from D. Sabatini (Massachusetts Institute of Technology, Cambridge, MA) via Addgene (Plasmid #14951, (Sancak et al., 2007)). Full length Rheb was PCR amplified from this vector and cloned into *Smal*-digested pEGFPC1 by Gibson Assembly (New England Biolabs, Ipswich, MA). The Q5 Site-Directed Mutagenesis Kit (New England Biolabs) was used to generate GFP-Rheb^ΔCAAX^, GFP-Rheb-HRas, and GFP-Rheb-HRas^C➔A^ plasmids. To generate GFP-Rheb-ER, a cDNA (IDT, Coralville, IA) containing a fragment of Cytochrome b5 (region from amino acids 95-134 (Itakura and Mizushima, 2010) with myc tag and glycosylation site (Honsho et al., 1998) was cloned into GFP-Rheb^ΔCAAX^ (linearized at the C Terminus via PCR) by Gibson Assembly. Full length RhebL1 cDNA was synthesized (IDT) and cloned into *Smal*-digested pEGFPC1 by Gibson Assembly. The following plasmids were kind gifts: LAMP1-mCherry from J. Lippincott-Schwartz (Janelia Research Campus, Ashburn, VA) and mRFP-Sec61B from T. Rapoport (Harvard University, Cambridge, MA). Oligonucleotide primers and cDNA sequences used to generate these plasmids are listed in Table S1.

Small guide RNAs (gRNAs) were designed using the CRISPR design tool (crispr.mit.edu) or selected from predesigned gRNA sequences (Sanjana et al., 2014). The gRNA-encoding DNA oligonucleotides (IDT) sequences were annealed and ligated into Bbs1-digested pX459 V2.0 plasmid that was provided by F. Zhang (Massachusetts Institute of Technology, Cambridge, MA) via Addgene (#62988, (Ran et al., 2013)) and transformed into Stabl3-competent *Escherichia coli* cells. The gRNA sequences are listed in Table S2.

### Immunoblotting

Cells were lysed in Tris-Buffered Saline + 1% Triton X-100 (TBST) with protease and phosphatase inhibitor cocktails (Roche Diagnostics, Florham Park, NJ). To remove insoluble materials, lysates were centrifuged for 6 minutes at 14,000 rpm. Lysate protein concentrations were measured via Bradford assay prior to denaturation with Laemmli buffer and 5-minutes at 95C. Immunoblotting was performed with 4-15% gradient Mini-PROTEAN TGX precast polyacrylamide gels and nitrocellulose membranes (Bio-Rad, Hercules, CA). Membranes were blocked with 5% milk in TBS with 0.1% Tween 20 (TBST) buffer and then incubated with antibodies in 5% milk or bovine serum albumin in TBST. Antibodies used in this study are summarized in the Table S4. Horseradish peroxidase signal detection was performed using chemiluminescent detection reagents (ThermoScientific) and the Versadoc imaging station (Bio-Rad). ImageJ (National Institutes of Health) was used to analyze the results and measure band intensities.

### Cell Fractionation

To perform cell fractionation, 2 million HeLa cells were plated in 10-cm dishes. Cells were rinsed, scraped into chilled PBS, and spun at 1000 rpm for 10 minutes. PBS was aspirated and cells were resuspended in homogenization buffer (HB; 5 mM Tris-HCl, 250 mM sucrose, 1 mM egtazic acid (EGTA), pH 7.4) with protease and phosphatase inhibitor cocktails. Cells were homogenized using a ball-bearing cell cracker (20 passages at 10 μm clearance, Isobiotec, Germany). Lysates were centrifuged at 1000 rpm for 10 minutes to remove unlysed cells. Supernatent was spun at 45,000 rpm for 1 hour to pellet membrane. Supernatant containing cytosolic proteins and the re-suspended membrane pellet were analyzed via immunoblotting.

### Immunofluorescence and Microscopy

Cells were grown on 12-mm No. 1 glass coverslips (Carolina Biological Supply) and fixed with 4% paraformaldehyde (PFA; Electron Microscopy Sciences, Hatfield, PA) in 0.1M sodium phosphate buffer (pH 7.2) for 30 minutes. To best preserve cell structure, this was achieved by adding 8% PFA dropwise to cells growing on coverslips in growth medium. Coverslips were washed in PBS before permeabilization in PBS + 0.2% Triton X-100 for 10 minutes. Coverslips were blocked for 30 minutes in blocking buffer (5% NDS/PBS/0.2% Triton X-100). Cells were incubated in primary antibody overnight at 4°C in blocking buffer. Cells were subsequently washed 3x with 0.2% Triton X-100 and incubated in secondary antibody in blocking buffer for thirty minutes at room temperature. Cells were washed 3x with 0.2% Triton X-100 before preparing slides with Prolong Gold Mounting media (Invitrogen, Carlsbad, CA). Antibodies used in this study are summarized in the Table S4.

For live cell imaging, cells were grown on glass bottom dishes (MatTek Corporation, Ashland, MA) prior to imaging. Sub-confluent dishes were imaged via spinning disc confocal microscopy at room temperature in a buffer that contained: 136 mM NaCl, 2.5 mM KCl, 2 mM CaCl_2_, 1.3 mM MgCl_2_ and 10 mM Hepes, 0.2% Glucose, 0.2% BSA pH 7.4 (Brown et al., 2000). Dextran Alexa Fluor 647 (Invitrogen, #22914) was applied overnight and washed out one hour before imaging. Our microscope consisted of the UltraVIEW VOX system (PerkinElmer, Waltham, MA) including the Ti-R Eclipse, Nikon inverted microscope (equipped with a 60x CFI Plan Apo VC, NA 1.4, oil immersion), a spinning disk confocal scan head (CSU-X1, Yokogawa) and Volocity (PerkinElmer) software. Images were acquired without binning with a 14-bit (1,000 x 1,000) EMCCD (Hamamatsu Photonics) and processed with ImageJ. ImageJ JACoP plug-in was used to generate correlation coefficients (Bolte and Cordelieres, 2006).

### CRISPR/Cas9 Genome Editing

Similar to a previously described protocol (Amick et al., 2018), to generate knock out cells, HeLa cells were co-transfected with Rheb and RhebL1 sgRNAs encoded within the pX459 plasmid. On the next day, puromycin (2 μg/ml) was added and cells were selected for 2 days. Surviving cells were plated at single-cell density and allowed to recover. Putative knockout colonies were initially identified via immunoblotting and subsequently confirmed by sequencing of the genomic loci. To obtain genomic DNA sequence for the target site in the Rheb gene, DNA was extracted (QuickExtract DNA extraction solution; Epicentre Biotechnologies), the region of interest was amplified by PCR (primers summarized in Table S3), PCR products were cloned into the pCR-Blunt TOPO vector (Zero Blunt TOPO PCR cloning kit; ThermoFisher Scientific), and transformed into XL-1 Blue Supercompetent Cells (Agilent Technologies, Santa Clara, CA). Plasmid DNA was isolated from multiple colonies and sequenced.

To insert a 2xHA epitope tag at the endogenous Rheb locus, we utilized a CRISPR/Cas9 genome-editing system strategy as previously described (Amick et al., 2016; Leonetti et al., 2016; Petit et al., 2013; Richardson et al., 2016). The crRNA, tracr RNA and ssDNA oligo were resuspended in nuclease-free duplex buffer (IDT) to 100 μM. To form the RNA duplex, the crRNA and tracrRNA were added at a 1:1 ratio and heated for 5 min at 95°C, then cooled to room temperature. The RNA duplex (150 pmol), Cas9 (150 pmol) and OptiMem were mixed in a sterile tube and incubated for 10 minutes at room temperature to form RNP complex. Hela cells (750,000 cells) were electroporated (Cell Line Kit V, electroporation program Q01; Lonza, Basel, Switzerland) with 10 μl RNP complex crRNA and single-stranded DNA (ssDNA) repair template (IDT Ultramer format). Sequences are provided in Table S2. Clonal cell lines were generated and validated using western blotting and immunofluorescence.

### Statistical Analysis

Data were analyzed using Prism (Graphpad software) and tests are denoted in figure legends. All error bars represent standard deviations. Data distribution was assumed to be normal, but this was not formally tested.

## References

Amick, J., A. Roczniak-Ferguson, and S.M. Ferguson. 2016. C9orf72 binds SMCR8, localizes to lysosomes, and regulates mTORC1 signaling. Mol Biol Cell. 27:3040–3051.

Amick, J., A.K. Tharkeshwar, C. Amaya, and S.M. Ferguson. 2018. WDR41 supports lysosomal response to changes in amino acid availability. Mol Biol Cell. 29:2213–2227.

Bar-Peled, L., L. Chantranupong, A.D. Cherniack, W.W. Chen, K.A. Ottina, B.C. Grabiner, E.D. Spear, S.L. Carter, M. Meyerson, and D.M. Sabatini. 2013. A Tumor suppressor complex with GAP activity for the Rag GTPases that signal amino acid sufficiency to mTORC1. Science. 340:1100–1106.

Bar-Peled, L., L.D. Schweitzer, R. Zoncu, and D.M. Sabatini. 2012. Ragulator is a GEF for the rag GTPases that signal amino acid levels to mTORC1. Cell. 150:1196–1208.

Ben-Sahra, I., and B.D. Manning. 2017. mTORC1 signaling and the metabolic control of cell growth. Curr Opin Cell Biol. 45:72–82.

Betz, C., and M.N. Hall. 2013. Where is mTOR and what is it doing there? J Cell Biol. 203:563–574.

Bolte, S., and F.P. Cordelieres. 2006. A guided tour into subcellular colocalization analysis in light microscopy. J Microsc. 224:213–232.

Brown, P.S., E. Wang, B. Aroeti, S.J. Chapin, K.E. Mostov, and K.W. Dunn. 2000. Definition of distinct compartments in polarized Madin-Darby canine kidney (MDCK) cells for membrane-volume sorting, polarized sorting and apical recycling. Traffic. 1:124–140.

Buerger, C., B. DeVries, and V. Stambolic. 2006. Localization of Rheb to the endomembrane is critical for its signaling function. Biochem Biophys Res Commun. 344:869–880.

Burd, C.G., and S.D. Emr. 1998. Phosphatidylinositol(3)-phosphate signaling mediated by specific binding to RING FYVE domains. Mol Cell. 2:157–162.

Cadwallader, K.A., H. Paterson, S.G. Macdonald, and J.F. Hancock. 1994. N-terminally myristoylated Ras proteins require palmitoylation or a polybasic domain for plasma membrane localization. Mol Cell Biol. 14:4722–4730.

Carroll, B., D. Maetzel, O.D. Maddocks, G. Otten, M. Ratcliff, G.R. Smith, E.A. Dunlop, J.F. Passos, O.R. Davies, R. Jaenisch, A.R. Tee, S. Sarkar, and V.I. Korolchuk. 2016. Control of TSC2-Rheb signaling axis by arginine regulates mTORC1 activity. Elife. 5.

Choy, E., V.K. Chiu, J. Silletti, M. Feoktistov, T. Morimoto, D. Michaelson, I.E. Ivanov, and M.R. Philips. 1999. Endomembrane trafficking of ras: the CAAX motif targets proteins to the ER and Golgi. Cell. 98:69–80.

Clark, G.J., M.S. Kinch, K. Rogers-Graham, S.M. Sebti, A.D. Hamilton, and C.J. Der. 1997. The Ras-related protein Rheb is farnesylated and antagonizes Ras signaling and transformation. J Biol Chem. 272:10608–10615.

Demetriades, C., N. Doumpas, and A.A. Teleman. 2014. Regulation of TORC1 in response to amino acid starvation via lysosomal recruitment of TSC2. Cell. 156:786–799.

Demetriades, C., M. Plescher, and A.A. Teleman. 2016. Lysosomal recruitment of TSC2 is a universal response to cellular stress. Nat Commun. 7:10662.

Ferguson, S.M. 2015. Beyond indigestion: emerging roles for lysosome-based signaling in human disease. Curr Opin Cell Biol. 35:59–68.

Gaullier, J.M., A. Simonsen, A. D’Arrigo, B. Bremnes, H. Stenmark, and R. Aasland. 1998. FYVE fingers bind PtdIns(3)P. Nature. 394:432–433.

Gonzalez, A., and M.N. Hall. 2017. Nutrient sensing and TOR signaling in yeast and mammals. EMBO J. 36:397–408.

Goorden, S.M., M. Hoogeveen-Westerveld, C. Cheng, G.M. van. Woerden, M. Mozaffari, L. Post, H.J. Duckers, M. Nellist, and Y. Elgersma. 2011. Rheb is essential for murine development. Mol Cell Biol. 31:1672–1678.

Groenewoud, M.J., S.M. Goorden, J. Kassies, W. Pellis-van Berkel, R.F. Lamb, Y. Elgersma, and F.J. Zwartkruis. 2013. Mammalian target of rapamycin complex I (mTORC1) activity in ras homologue enriched in brain (Rheb)-deficient mouse embryonic fibroblasts. PLoS One. 8:e81649.

Hancock, J.F., H. Paterson, and C.J. Marshall. 1990. A polybasic domain or palmitoylation is required in addition to the CAAX motif to localize p21ras to the plasma membrane. Cell. 63:133–139.

Hanker, A.B., N. Mitin, R.S. Wilder, E.P. Henske, F. Tamanoi, A.D. Cox, and C.J. Der. 2010. Differential requirement of CAAX-mediated posttranslational processing for Rheb localization and signaling. Oncogene. 29:380–391.

Hao, F., K. Kondo, T. Itoh, S. Ikari, S. Nada, M. Okada, and T. Noda. 2018. Rheb localized on the Golgi membrane activates lysosome-localized mTORC1 at the Golgi-lysosome contact site. J Cell Sci. 131.

Honsho, M., J.Y. Mitoma, and A. Ito. 1998. Retention of cytochrome b5 in the endoplasmic reticulum is transmembrane and luminal domain-dependent. J Biol Chem. 273:20860–20866.

Ismail, S.A., Y.X. Chen, A. Rusinova, A. Chandra, M. Bierbaum, L. Gremer, G. Triola, H. Waldmann, P.I. Bastiaens, and A. Wittinghofer. 2011. Arl2-GTP and Arl3-GTP regulate a GDI-like transport system for farnesylated cargo. Nat Chem Biol. 7:942–949.

Itakura, E., and N. Mizushima. 2010. Characterization of autophagosome formation site by a hierarchical analysis of mammalian Atg proteins. Autophagy. 6:764–776.

Kovacevic, M., C.H. Klein, L. Rossmannek, A.D. Konitsiotis, A. Stanoev, A.U. Kraemer, and P.I. Bastiaens. 2018. A spatially regulated GTPase cycle of Rheb 1 controls growth factor signaling to mTORC1. BioRXiv.

Leonetti, M.D., S. Sekine, D. Kamiyama, J.S. Weissman, and B. Huang. 2016. A scalable strategy for high-throughput GFP tagging of endogenous human proteins. Proc Natl Acad Sci U S A. 113:E3501–3508.

Lim, C.Y., and R. Zoncu. 2016. The lysosome as a command-and-control center for cellular metabolism. J Cell Biol. 214:653–664.

Menon, S., C.C. Dibble, G. Talbott, G. Hoxhaj, A.J. Valvezan, H. Takahashi, L.C. Cantley, and B.D. Manning. 2014. Spatial control of the TSC complex integrates insulin and nutrient regulation of mTORC1 at the lysosome. Cell. 156:771–785.

Papke, B., S. Murarka, H.A. Vogel, P. Martin-Gago, M. Kovacevic, D.C. Truxius, E.K. Fansa, S. Ismail, G. Zimmermann, K. Heinelt, C. Schultz-Fademrecht, A. Al Saabi, M. Baumann, P. Nussbaumer, A. Wittinghofer, H. Waldmann, and P.I. Bastiaens. 2016. Identification of pyrazolopyridazinones as PDEdelta inhibitors. Nat Commun. 7:11360.

Petit, C.S., A. Roczniak-Ferguson, and S.M. Ferguson. 2013. Recruitment of folliculin to lysosomes supports the amino acid-dependent activation of Rag GTPases. J Cell Biol. 202:1107–1122.

Ran, F.A., P.D. Hsu, J. Wright, V. Agarwala, D.A. Scott, and F. Zhang. 2013. Genome engineering using the CRISPR-Cas9 system. Nat Protoc. 8:2281–2308.

Richardson, C.D., G.J. Ray, M.A. DeWitt, G.L. Curie, and J.E. Corn. 2016. Enhancing homology-directed genome editing by catalytically active and inactive CRISPR-Cas9 using asymmetric donor DNA. Nat Biotechnol. 34:339–344.

Rocks, O., A. Peyker, M. Kahms, P.J. Verveer, C. Koerner, M. Lumbierres, J. Kuhlmann, H. Waldmann, A. Wittinghofer, and P.I. Bastiaens. 2005. An acylation cycle regulates localization and activity of palmitoylated Ras isoforms. Science. 307:1746–1752.

Rowland, A.A., P.J. Chitwood, M.J. Phillips, and G.K. Voeltz. 2014. ER contact sites define the position and timing of endosome fission. Cell. 159:1027–1041.

Sancak, Y., L. Bar-Peled, R. Zoncu, A.L. Markhard, S. Nada, and D.M. Sabatini. 2010. Ragulator-Rag complex targets mTORC1 to the lysosomal surface and is necessary for its activation by amino acids. Cell. 141:290–303.

Sancak, Y., T.R. Peterson, Y.D. Shaul, R.A. Lindquist, C.C. Thoreen, L. Bar-Peled, and D.M. Sabatini. 2008. The Rag GTPases bind raptor and mediate amino acid signaling to mTORC1. Science. 320:1496–1501.

Sancak, Y., C.C. Thoreen, T.R. Peterson, R.A. Lindquist, S.A. Kang, E. Spooner, S.A. Carr, and D.M. Sabatini. 2007. PRAS40 is an insulin-regulated inhibitor of the mTORC1 protein kinase. Mol Cell. 25:903–915.

Sanjana, N.E., O. Shalem, and F. Zhang. 2014. Improved vectors and genome-wide libraries for CRISPR screening. Nat Methods. 11:783–784.

Sato, T., A. Nakashima, L. Guo, and F. Tamanoi. 2009. Specific activation of mTORC1 by Rheb G-protein in vitro involves enhanced recruitment of its substrate protein. J Biol Chem. 284:12783–12791.

Saxton, R.A., and D.M. Sabatini. 2017. mTOR Signaling in Growth, Metabolism, and Disease. Cell. 168:960–976.

Silvius, J.R., P. Bhagatji, R. Leventis, and D. Terrone. 2006. K-ras4B and prenylated proteins lacking “second signals” associate dynamically with cellular membranes. Mol Biol Cell. 17:192–202.

Silvius, J.R., and F. l’Heureux. 1994. Fluorimetric evaluation of the affinities of isoprenylated peptides for lipid bilayers. Biochemistry. 33:3014–3022.

Smeland, T.E., M.C. Seabra, J.L. Goldstein, and M.S. Brown. 1994. Geranylgeranylated Rab proteins terminating in Cys-Ala-Cys, but not Cys-Cys, are carboxyl-methylated by bovine brain membranes in vitro. Proc Natl Acad Sci US A. 91:10712–10716.

Takahashi, K., M. Nakagawa, S.G. Young, and S. Yamanaka. 2005. Differential membrane localization of ERas and Rheb, two Ras-related proteins involved in the phosphatidylinositol 3-kinase/mTOR pathway. J Biol Chem. 280:32768–32774.

Tee, A.R., J. Blenis, and C.G. Proud. 2005. Analysis of mTOR signaling by the small G-proteins, Rheb and RhebL1. FEBS Lett. 579:4763–4768.

Tsun, Z.Y., L. Bar-Peled, L. Chantranupong, R. Zoncu, T. Wang, C. Kim, E. Spooner, and D.M. Sabatini. 2013. The folliculin tumor suppressor is a GAP for the RagC/D GTPases that signal amino acid levels to mTORC1. Mol Cell. 52:495–505.

Walton, Z.E., C.H. Patel, R.C. Brooks, Y. Yu, A. Ibrahim-Hashim, M. Riddle, A. Porcu, T. Jiang, B.L. Ecker, F. Tameire, C. Koumenis, A.T. Weeraratna, D.K. Welsh, R. Gillies, J.C. Alwine, L. Zhang, J.D. Powell, and C.V. Dang. 2018. Acid Suspends the Circadian Clock in Hypoxia through Inhibition of mTOR. Cell. 174:72–87 e32.

Wolfson, R.L., L. Chantranupong, G.A. Wyant, X. Gu, J.M. Orozco, K. Shen, K.J. Condon, S. Petri, J. Kedir, S.M. Scaria, M. Abu-Remaileh, W.N. Frankel, and D.M. Sabatini. 2017. KICSTOR recruits GATOR1 to the lysosome and is necessary for nutrients to regulate mTORC1. Nature. 543:438–442.

Yang, H., X. Jiang, B. Li, H.J. Yang, M. Miller, A. Yang, A. Dhar, and N.P. Pavletich. 2017. Mechanisms of mTORC1 activation by RHEB and inhibition by PRAS40. Nature. 552:368–373.

Zahonova, K., R. Petrzelkova, M. Valach, E. Yazaki, D.V. Tikhonenkov, A. Butenko, J. Janouskovec, S. Hrda, V. Klimes, G. Burger, Y. Inagaki, P.J. Keeling, V. Hampl, P. Flegontov, V. Yurchenko, and M. Elias. 2018. Extensive molecular tinkering in the evolution of the membrane attachment mode of the Rheb GTPase. Sci Rep. 8:5239.

Zimmermann, G., B. Papke, S. Ismail, N. Vartak, A. Chandra, M. Hoffmann, S.A. Hahn, G. Triola, A. Wittinghofer, P.I. Bastiaens, and H. Waldmann. 2013. Small molecule inhibition of the KRAS-PDEdelta interaction impairs oncogenic KRAS signalling. Nature. 497:638–642.

