## Supplemental Data for "Weak Membrane Interactions Allow Rheb to Activate mTORC1 Signaling Without Major Lysosome Enrichment"

Supplemental Figure 1

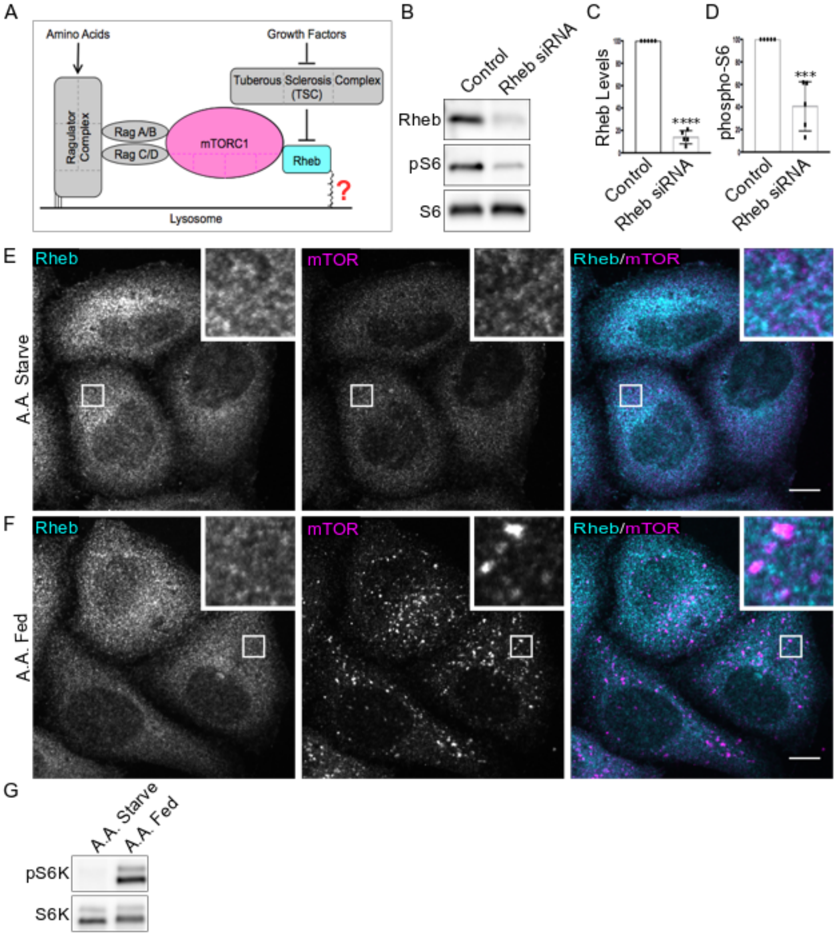


**Supplemental Figure 1. Subcellular localization of Rheb is not responsive to changes in amino acid availability. (A)** Schematic diagram depicting the widely accepted model for mTORC1 activation on the surface of lysosomes. **(B)** Immunoblot analysis of Control and Rheb siRNA-treated HeLa cells. **(C)** Quantification of Rheb protein levels in Control and Rheb siRNA-treated cells (****, P<0.0001; unpaired t-test; n=5). **(D)** Quantification of phospho-S6 levels in Control and Rheb siRNA-treated cells (***, P=0.0003; unpaired t-test; n=5). **(E)** Representative immunofluorescence images of Rheb and mTOR localization in HeLa cells after a 2-hour amino acid (A.A) starvation. **(F)** Representative immunofluorescence images of mTOR and Rheb localization in HeLa cells following 20-minute A.A. refeeding of starved cells. **(G)** Immunoblot analysis of phospho-S6K and S6K levels in starved and re-fed HeLa cells. Scale bars, 10 µm.

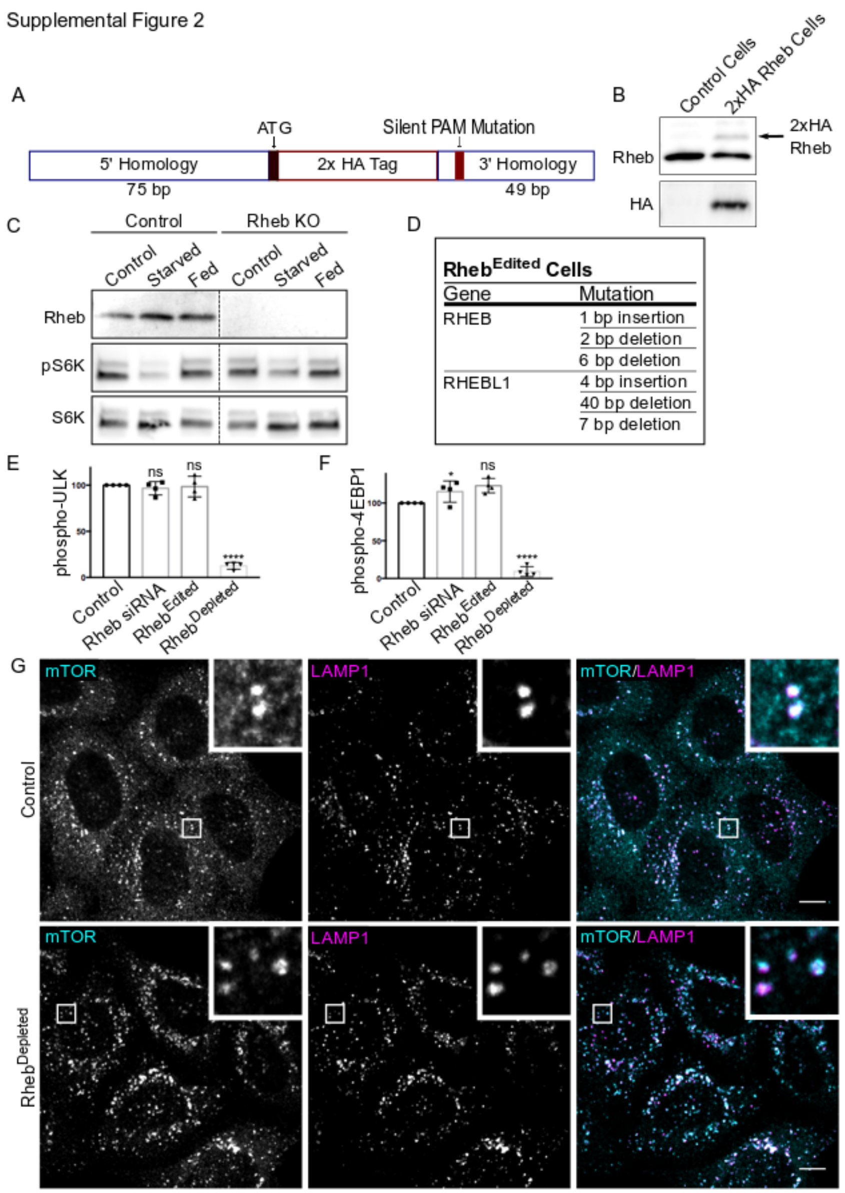


**Supplemental Figure 2. Characterization of CRISPR-edited Rheb and RhebL1 mutant cells.** **(A)** Summary of the genome editing strategy to used to insert a 2xHA epitope tag into the endogenous Rheb locus. **(B)** Western blot analysis of Control HeLa cells and 2xHA Rheb HeLa cells with anti-HA and anti-Rheb antibodies. **(C)** Immunoblot analysis of Rheb and phosphorylated S6K levels in CRISPR-mediated Rheb single knock out (KO) cells. Control and KO cells were starved of growth factors overnight and re-fed with serum for 30 minutes. **(D)** Summary of CRISPR-Cas9 mediated indels in $Rheb^{Edited}$/ $Rheb^{Depleted}$ cells identified by sequencing of genomic DNA. **(E)** Quantification of phospho-Ulk in Control and Rheb siRNA-treated, $Rheb^{Edited}$, and $Rheb^{Depleted}$HeLa cells (from Figure 3A). The phospho-Ulk levels were divided by Ulk and normalized to Control (****, P<0.0001; ANOVA with Dunnett's Multiple Comparisons Test, n=4). **(F)** Quantification of phospho-4EBP1 in Control and Rheb siRNA-treated, $Rheb^{Edited}$, and $Rheb^{Depleted}$HeLa cells (from Figure 3A). phospho-4EBP1 levels were divided by 4EBP1 and normalized to Control (*, P<0.05; ****, P<0.0001; ANOVA with Dunnett's Multiple Comparisons Test, n=4). **(G)** Immunofluorescence images of the subcellular localization of mTOR and LAMP1 in control and $Rheb^{Depleted}$HeLa cells. Scale bars, 10 μm.

Supplemental Figure 3

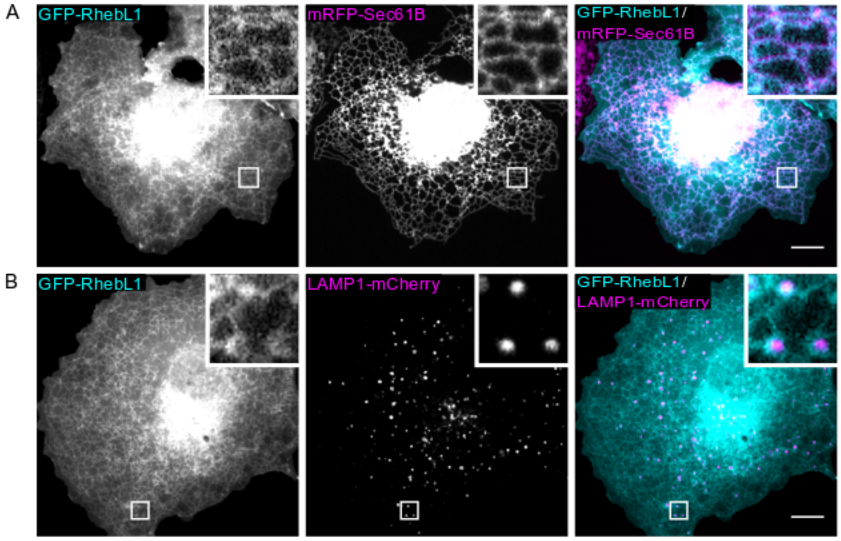


**Supplemental Figure 3. GFP-RhebL1 localizes to ER and Cytosol. (A)** GFP-RhebL1and mRFP-Sec61B sub-cellular localization in COS-7 cells. **(B)** GFP-RhebL1 and LAMP1-mCherry sub-cellular localization. Scale bars, 10 μm.

Supplemental Figure 4

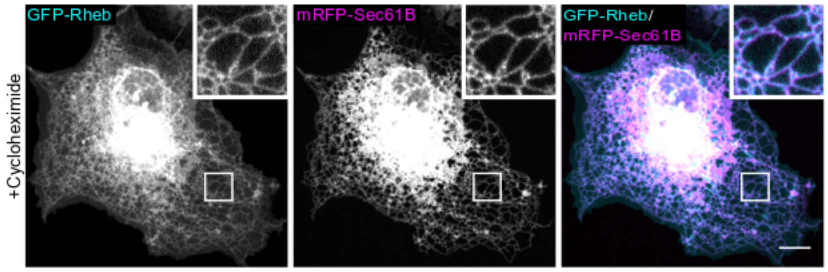


**Supplemental Figure 4. ER localization of GFP-Rheb is not sensitive to inhibition of new protein synthesis.** Live-cell imaging of GFP-Rheb localization following a 2 hour treatment of COS-7 cells with cycloheximide (100ug/mL). Scale bar, 10 μm.

**Supplemental Movie 1. GFP-Rheb localizes to dynamic ER tubules that contact lysosomes.** Spinning disk confocal live-cell imaging of GFP-Rheb (cyan) and Dextran-loaded lysosomes (magenta; cells were labeled overnight with Alexa647-Dextran and washed for 1 hour before imaging). Images were acquired at a rate of 15 frames/minute over an interval of 5 minutes. Scale bar, 5 µm.

**Table S1: Summary of Oligonucleotides used in Plasmid Construction**

| Target | Use | Sequence (5'-3') |
|---|---|---|
| Rheb Forward | Rheb forward primer for cloning into pEGFP-C1 | **CAGTCGACGGTACCGCGGGCCC**aATGCCGCAGTCCAAGTCCCGGA |
| Rheb Reverse | Rheb reverse primer for cloning into into pEGFP-C1 | **CAGTTATCTAGATCCGGTGGATCCC**TCACATCACCGAGCACGAA |
| Rheb$^{CaaX}$ Forward | Site directed mutagenesis (SDM) forward primer to delete Rheb CaaX box | TGAGGGATCCACCGGATC |
| Rheb$^{CaaX}$ Reverse | SDM reverse primer to delete Rheb CaaX box | CGAAGACTTCCCTTGTGAAG |
| Rheb-H-Ras Forward | SDM forward primer to replace Rheb hypervariable region (HVR) with HRas HVR | ctgcatgagctgcaagtgtgtgctctcctgaGGGATCCACCGGATCTAG |
| Rheb-H-Ras Reverse | SDM reverse primer to replace Rheb HVR with H-Ras HVR | ccggggccactctcatcaggagggttcagcttTTTTTCTGCCTCCAAAATTATCC |
| Rheb-H-Ras$^{C\rightarrow A}$ Forward | SDM forward primer to mutate C to A in H-Ras HVR | agcgccAAGTGTGTGCTCTCCTGAGC |
| Rheb-H-Ras $^{C\rightarrow A}$ Reverse | SDM reverse primer to mutate C to A in H-Ras HVR | catggcGCCGGGGCCACTCTCATC |
| Myr-Rheb$^{CaaX}$ Forward | SDM forward primer to add myristoylation consensus sequence to Rheb$^{CaaX}$ | cggaggatccggcggaCCGCAGTCCAAGTCCCGG |
| Myr-Rheb$^{CaaX}$ Reverse | SDM reverse primer to add myristoylation consensus sequence to Rheb$^{CaaX}$ | gatccgccggatcctccTGTAAGAGATTGACCCATGGTGG |
| NonMyr-Rheb$^{CaaX}$ Forward | SDM forward primer to add mutated myristoylation sequence to Rheb$^{CaaX}$ | ggcggatccggaggatccggcggaCCGCAGTCCAAGTCCCGG |
| NonMyr-Rheb$^{CaaX}$ Reverse | SDM reverse primer to add mutated myristoylation sequence to Rheb$^{CaaX}$ | ggatcctcctgtaagagattgggcCATGGTGGCGACCGGTAG |
| Cytochrome b5 cDNA | cDNA added to C Terminus of Rheb$^{CaaX}$ to generate GFP-Rheb-ER | AGCTTCACAAGGGAAGTCTTCGccggaaactcttatcactactattgattctagttccagttggtggaccaactgggtgatccctgccatctctgcagtggccgtcgccttgatgtatcgcctatacatggcagaggactgaGGGATCCACCGGATCTAGATAA |
| RhebL1 cDNA | cDNA with overhangs to insert RhebL1 into pEGFP-C1 | GCAGTCGACGGTACCGCGGGCCCaATGCCGCTAGTCCGCTACAGGAAGGTGGTCATCCTCGGATACCGCTGTGTAGGGAAGACATCTTTGGCACATCAATTTGTGGAAGGCGAGTTCTCGGAAGGCTACGATCCTACAGTGGAGAATACTTACAGCAAGATAGTGACTCTTGGCAAAGATGAGTTTCACCTACATCTGGTGGACACAGCAGGGCAGGATGAGTACAGCATTCTGCCCTATTCATTCATCATTGGGGTCCATGGTTATGTGCTTGTGTATTCTGTCACCTCTCTGCATAGCTTC |

CAAGTCATTGAGAGTCTGTACCAAAAGC
TACATGAAGGCCATGGGAAAACCCGGG
TGCCAGTGGTTCTAGTGGGGAACAAGG
CAGATCTCTCTCCAGAGAGAGAGGTAC
AGGCAGTTGAAGGAAAGAAGCTGGCAG
AGTCCTGGGGTGCGACATTTATGGAGT
CATCTGCTCGAGAGAATCAGCTGACTCA
AGGCATCTTCACCAAAGTCATCCAGGA
GATTGCCCGTGTGGAGAATTCCTATGG
GCAAGAGCGTCGCTGCCATCTCATGTG
AGGGATCCACCGGATCTAGATAAC

**Table S2: Summary of Oligonucleotides for Gene-Editing Experiments**

| Gene | Use | Sense (5'-3') | Antisense (5'-3') |
|---|---|---|---|
| RHEB | gRNA for KO | CACCGCTACATACCTTTCCATATGC | AAACGCATATGGAAAGGTATGTAGC |
| RHEBL1 | gRNA for KO | CACCGATCCTCGGATACCGCTGTGT | AAACACACAGCGGTATCCGAGGATC |
| RHEB | crRNA for 2xHA tag insertion | GAUCGCGAUCUUCCGGGACUGUUUUAGAGCUAUGCU | |
| RHEB | 2xHA tag template | GCCGCCGATCACAGCAGCAGGAGCCACCGCCGCCGCGGTTGATGTGGTTGGGCCGGGGCTGAGGAGGCCGCCAAGATGtacccatacgatgttccagattacgcttatccctatgacgtcccggactatgcaCCGCAGTCTAAGTCCCGGAAGATCGCGATCCTGGGCTACCGGTCTGTGG | |

**Table S3: Summary of Oligonucleotide Primers for PCR Amplification of Genomic DNA**

| Target | Sense (5'-3') | Antisense (5'-3') |
|---|---|---|
| RHEB | GTATCTTCTGAGCAATACAATC | ACCATAACACCTCTGAACGGAT |
| RHEBL1 | ACTTTCCTGCACCAGCTGCCG | TTCCAATCCTCGCTCTACAAC |

**Table S4: Summary of Antibodies**

| Target Protein | Source | Catalog No. |
|---|---|---|
| 4EBP1 | Cell Signaling Technology | 9644S |
| P-4EBP1 (T37/46) | Cell Signaling Technology | 2855S |
| Anti-Rabbit, HRP | Cell Signaling Technology | 7074S |
| Anti-Mouse, HRP | Cell Signaling Technology | 7076S |
| Anti-biotin, HRP | Cell Signaling Technology | 7075P5 |
| Calnexin | Cell Signaling Technology | 2679S |
| Donkey anti-Mouse, Alexa Fluor 488 | Thermo Fischer Scientific | A21202 |
| Donkey anti-Mouse, Alexa Fluor 594 | Thermo Fischer Scientific | A21203 |
| Donkey anti-Rabbit, Alexa Fluor 488 | Thermo Fischer Scientific | A21206 |
| Donkey anti-Mouse, Alexa Fluor 594 | Thermo Fischer Scientific | A21207 |
| GFP-HRP | Rockland | 600-103-215 |
| HA | Cell Signaling Technology | 2367S |
| HA-HRP | Roche | 12 013 819 001 |
| LAMP1 | DSHB | H4A3 |
| LAMP1 | Cell Signaling Technology | 9091S |
| mTOR | Cell Signaling Technology | 2983S |
| p70 S6K | Cell Signaling Technology | 9202S |
| P-p70 S6K (T389) | Cell Signaling Technology | 9234S |
| Rheb | Abnova | H00006009-M01 |
| S6 | Cell Signaling Technology | 2217S |
| P-S6 (S235/236) | Cell Signaling Technology | 4858S |
| ULK | Cell Signaling Technology | 8054S |
| P-ULK (S757) | Cell Signaling Technology | 14202S |
